# Substrate Specificity for Human Histidine Methyltransferase SETD3

**DOI:** 10.1101/2021.12.30.474520

**Authors:** Jordi C. J. Hintzen, Huida Ma, Hao Deng, Apolonia Witecka, Steffen B. Andersen, Jakub Drozak, Hong Guo, Ping Qian, Haitao Li, Jasmin Mecinović

## Abstract

Histidine methyltransferase SETD3 plays an important role in human biology and diseases. Previously, we showed that SETD3 catalyzes N^3^-methylation of histidine 73 in β-actin (Kwiatkowski et al., 2018). Here we report integrated synthetic, biocatalytic, biostructural and computational analyses on human SETD3-catalyzed methylation of β-actin peptides possessing histidine and its structurally and chemically diverse mimics. Our enzyme assays supported by biostructural analyses demonstrate that SETD3 has a broader substrate scope beyond histidine, including N-nucleophiles on the aromatic and aliphatic side chains. Quantum mechanical/molecular mechanical (QM/MM) molecular dynamics and free-energy simulations provide insight into binding geometries and the free energy barrier for the enzymatic methyl transfer to histidine mimics, further supporting experimental data that histidine is the superior SETD3 substrate over its analogs. This work demonstrates that human SETD3 has a potential to catalyze efficient methylation of several histidine mimics, overall providing mechanistic, biocatalytic and functional insight into β-actin histidine methylation by SETD3.

## Introduction

Actin is an essential constituent of microfilaments that form the cytoskeleton of eukaryotic cells, and thus plays an important role in maintaining the structure of the cell (dos Remedios et al., 2003; Dominguez and Holmes, 2011; Pollard and Cooper, 2009; Campellone and Welch, 2010). Human actin exists in six isoforms that show different cellular expression patterns and functions (Perrin and Ervasti, 2010). Among these six isoforms, β-actin (βA) is expressed ubiquitously in the cell and plays the central role in cytoskeletal stability and cell mobility (Letterrier et al., 2017). Actin exists freely in the cell as a monomeric globular protein (G-actin), which upon binding to ATP polymerizes and forms stable actin filaments (F-actin) (Pieters et al., 2016). These filaments represent a crucial part of the cellular architecture, and their length is controlled by hydrolysis of the terminal phosphate of bound ATP, which is released slowly and leads to depolymerization of the filament (Lappalainen, 2016; Varland et al., 2019). To fine-tune their physiological roles, many posttranslational modifications (PTMs) – including methylation, acetylation, SUMOylation and ubiquitination – have been found on actin proteins (Terman and Kashina, 2013). N^3^-methylation of the histidine residue at position 73 in βA (βA-His73) has been shown to be conserved among eukaryotic actin isoforms, and leads to a decreased rate of hydrolysis of the actin-bound ATP (Kabsch et al., 1990; Nyman et al., 2002). Histidine methylation prevents primary dystocia and plays a key role in parturition in female mice (Wilkinson et al., 2019). SETD3 was recently identified as the actin-specific histidine N-methyltransferase that in the presence of the S-adenosylmethionine (SAM) cosubstrate catalyzes N^3^-methylation of His73 in βA (Figure 1a) (Wilkinson et al., 2019; Kwiatkowski et al., 2018; Witecka et al., 2021). SETD3 is a member of the SET-domain containing methyltransferases, a large family of enzymes that catalyze methylation of lysine and arginine residues on histones and non-histone proteins (Carlson and Gozani, 2016). The SET-domain is an ∼130 amino acids long motif, which stands for **S**u(var)3-9, **E**nhancer of zeste and **T**rithorax (Dillon et al., 2005). Initially, SETD3 was identified as a histone lysine methyltransferase that catalyzes mono- and dimethylation of histone H3 at positions K4 and K36 (Eom et al., 2011; Wagner and Carpenter, 2012), however, a H3 peptide was shown to be a very poor substrate for SETD3 in comparison to a β-actin peptide fragment (Guo et al., 2019).

**Figure 1.**
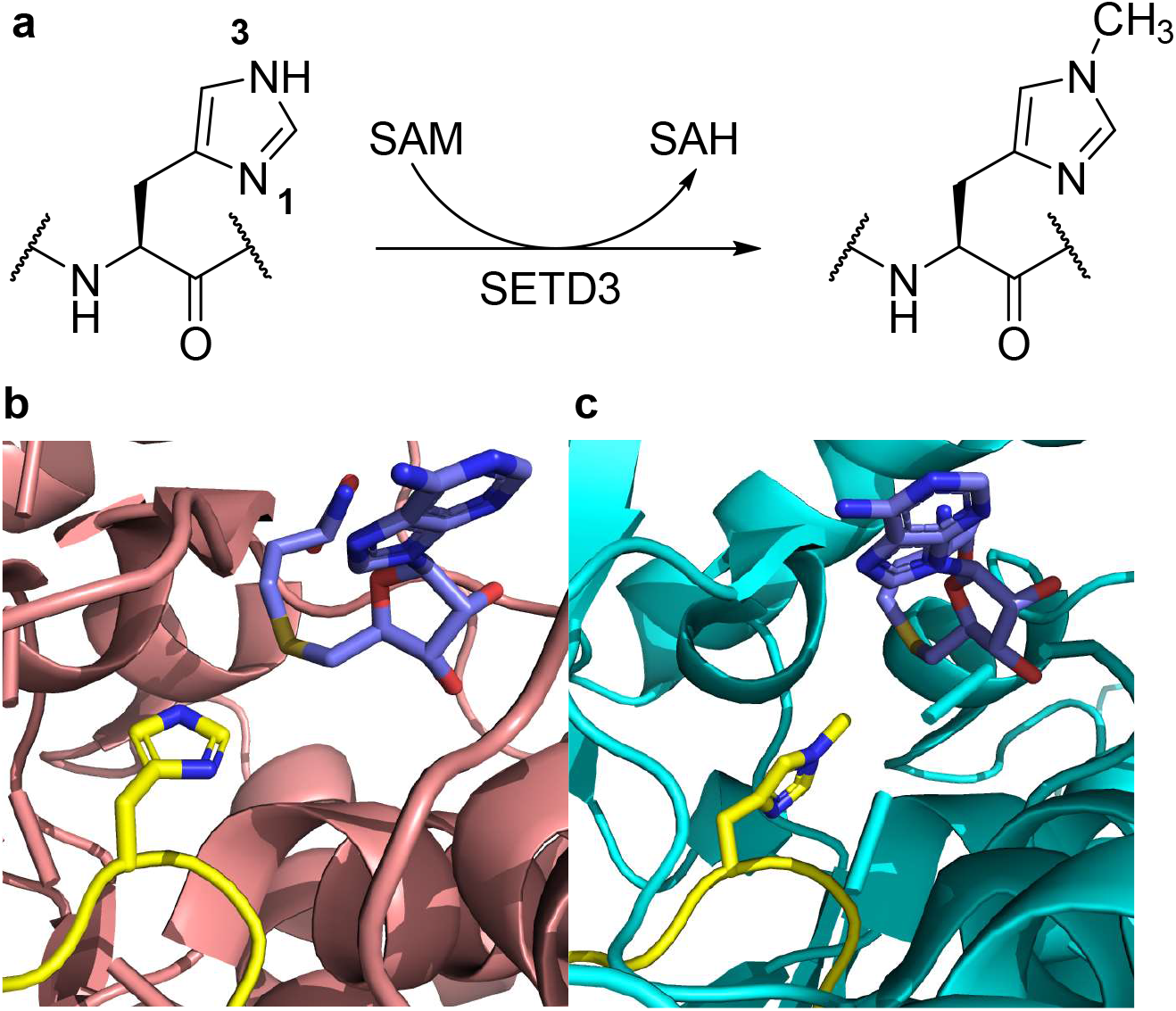
**a)** SETD3-catalyzed N^3^-methylation of histidine in βA, **b)** View from a crystal structure of SETD3 (pink) complexed with an unmodified βA peptide (yellow) and SAH, an unreactive product of the methylation reaction (blue) (PDB ID: 6ICV), **c)** View from a crystal structure of SETD3 (cyan) complexed with a methylated βA peptide (yellow) and SAH (blue) (PDB ID: 6ICT).

As the imidazole ring of histidine contains two nitrogen atoms at the N^1^ and N^3^ position (it is noteworthy that there has been often a confusion in the literature about the numbering of N atoms in histidine, they are also called N_τ1_ and N_ε2_ respectively) (Figure 1a), SETD3 could in principle have an ability to catalyze methylation of either or even both of these nitrogens. X-ray crystallographic and biochemical studies have shown, however, that SETD3 specifically catalyzes the methylation of histidine at the N^3^ position, *i*.*e*. a more distant nitrogen atom at the histidine side chain (Figure 1a) (Wilkinson et al., 2019, Guo et al., 2019, Dai et al., 2019, Zheng et al., 2020). Structural analyses of ternary complexes revealed that the βA peptide lies in a narrow groove formed by the SET domain of SETD3, with the imidazole ring of His73 buried in a hydrophobic pocket (Figure 1b). In the crystal structures containing the unmethylated His73 and *S*-adenosylmethionine (SAH) (Figure 1b), N^3^ of His73 in the βA peptide donates a hydrogen bond to the backbone carbonyl group of Asp275 and therefore exists as the N^3^-H τ tautomeric form that cannot undergo the methylation. Two different mechanisms seem to be possible. One is that there would be some general acid/base catalyst(s) that covert the N^3^-H τ tautomer to the N^1^-H π tautomer for methylation (Guo et al., 2019; Dai et al., 2019; Deng et al., 2020). Nevertheless, such general acid/base catalysts have not been identified with clarity, and replacement of some potential residues at the active site, for instance Asn255 by Ala or Tyr312 by Phe, did not abolish the enzyme’s activity. Alternatively, it was proposed that His73 in the N^1^-H π tautomeric form may bind to SETD3 directly for methylation (Deng et al., 2020). This suggestion seems to be consistent with several of the available crystal structures for actin (Nair et al., 2008, Rould et al., 2006), which show that the N^1^ atom of His73 (or from His73Me after the methylation on N^3^) donates a hydrogen bond to the backbone carbonyl oxygen of Gly158, indicating His73 of actin may exist as the N^1^-H π tautomer. In either of the two mechanisms, the chemical process of methylation is expected to start from the SETD3, N^1^-H π tautomer complex. Computer simulations demonstrated that the N^1^-H π tautomer was stable and well positioned and could undergo methylation at the SETD3 active site (Deng et al., 2020). Interestingly, the imidazole ring of the N^1^-H π tautomer from the simulations was found to have an orientation that is similar to the one in the crystal structures of the product complex containing His73Me (Figure 1c), suggesting that the imidazole ring might not undergo rotation during the methylation reaction.

Recent biochemical and biostructural studies have revealed that SETD3 exhibits a selectivity for histidine over lysine and methionine (Guo et al., 2019, Dai et al., 2019, Dai et. al. 2020a, Dai et al., 2020b). The histidine-binding pocket appears to be too small to efficiently accommodate the longer aliphatic side chain of lysine, although lysine’s side chain could be traced around the outer part of histidine’s imidazole ring, leading to the lysine’s terminal amine positioned alike the N^3^ of histidine’s imidazole. However, it was found that a crucial tyrosine residue, which forms a cation–π interaction with lysine in SET domain-containing histone lysine methyltransferases (Qian and Zhou, 2006), is not present in SETD3 where it is replaced by asparagine (Asn255) that seems to play an essential role in substrate binding and histidine methylation, forming a stabilizing hydrogen bond with the N^1^-H π tautomer of histidine (Deng et al., 2020) or the methylated histidine in product. Further cementing SETD3 as a histidine specific methyltransferase, an engineered SETD3 was generated wherein Asn255 was replaced by Phe, as well as Trp273 substituted by Ala to allow for more physical space for lysine’s side chain and to prevent steric clash in the hydrophobic binding pocket; these modifications caused SETD3 to display a 13-fold preference for lysine over histidine in a βA peptide (Dai et al., 2020b). Furthermore, SETD3 was also shown to have an ability to poorly methylate methionine generating *S*-methylmethionine (Dai et al., 2020a). Interestingly, the methionine containing βA peptide has a 76-fold stronger binding affinity and efficiently inhibits the activity of SETD3 (Dai et al., 2020a). βA peptides possessing methionine analogs at the His73 position were shown to inhibit SETD3 with nanomolar potency (Hintzen et al., 2021). Owing to SETD3’s unique reactivity towards histidine over lysine and methionine we sought to explore the substrate specificity of SETD3 to gain a deeper understanding of the underlying mechanism and biocatalytic scope of the SETD3 histidine methyltransferase. Here we report biomolecular studies on human SETD3-catalyzed methylation of histidine and its simplest mimics incorporated into βA peptides. Our synergistic synthetic, enzymatic, biostructural and computational work demonstrates that SETD3 has a broader substrate specificity beyond histidine.

## Results and Discussion

### The panel of histidine analogs

To investigate SETD3’s unique ability to catalyze an efficient methylation of histidine, we set out to incorporate a structurally and chemically diverse panel of histidine analogs into the βA peptide and examine the substrate scope of the human SETD3-catalyzed methylation reactions of βA (Figure 2). We have designed a large panel of histidine analogs that cover a wide chemical space, thus advancing basic understanding of SETD3 catalysis and its biocatalytic potential. The following histidine analogs have been selected: i) D-histidine to probe the stereochemistry; ii) N_α_-methylhistidine to explore the importance of the backbone; iii) premethylated N^1^-methyl-histidine and N^3^-methyl-histidine to further confirm the site of methylation; iv) C^2^-methyl-histidine to probe for steric effect on the imidazole ring; v) two triazolyl alanines and the tetrazolyl alanine to probe the most subtle change in reactivity of the imidazole ring; vi) pyridyl alanines and *para*-amino phenylalanine to investigate both the steric effects of a six-membered ring and the subtle change in the nitrogen’s location in the catalytic pocket of SETD3; vii) 3-(4-thiazolyl) alanine and tyrosine to explore methylation of sulfur or oxygen nucleophiles, respectively, and viii) lysine analogs to establish the optimal side chain length for SETD3-catalyzed methylation of aliphatic amines. All histidine analogs were incorporated at position 73 of the synthetic βA peptide (16-mer, residues 66-81) using automated, microwave assisted solid-phase peptide synthesis (SPPS). Coupling of the Fmoc-protected histidine analogs went smoothly, with no indicative change in reactivity observed in comparison to L-histidine. After cleavage from the Rink-Amide resin with TFA, all synthetic βA peptides were purified by reverse phase HPLC and lyophilized (Table S1, Figures S1-20).

**Figure 2.**
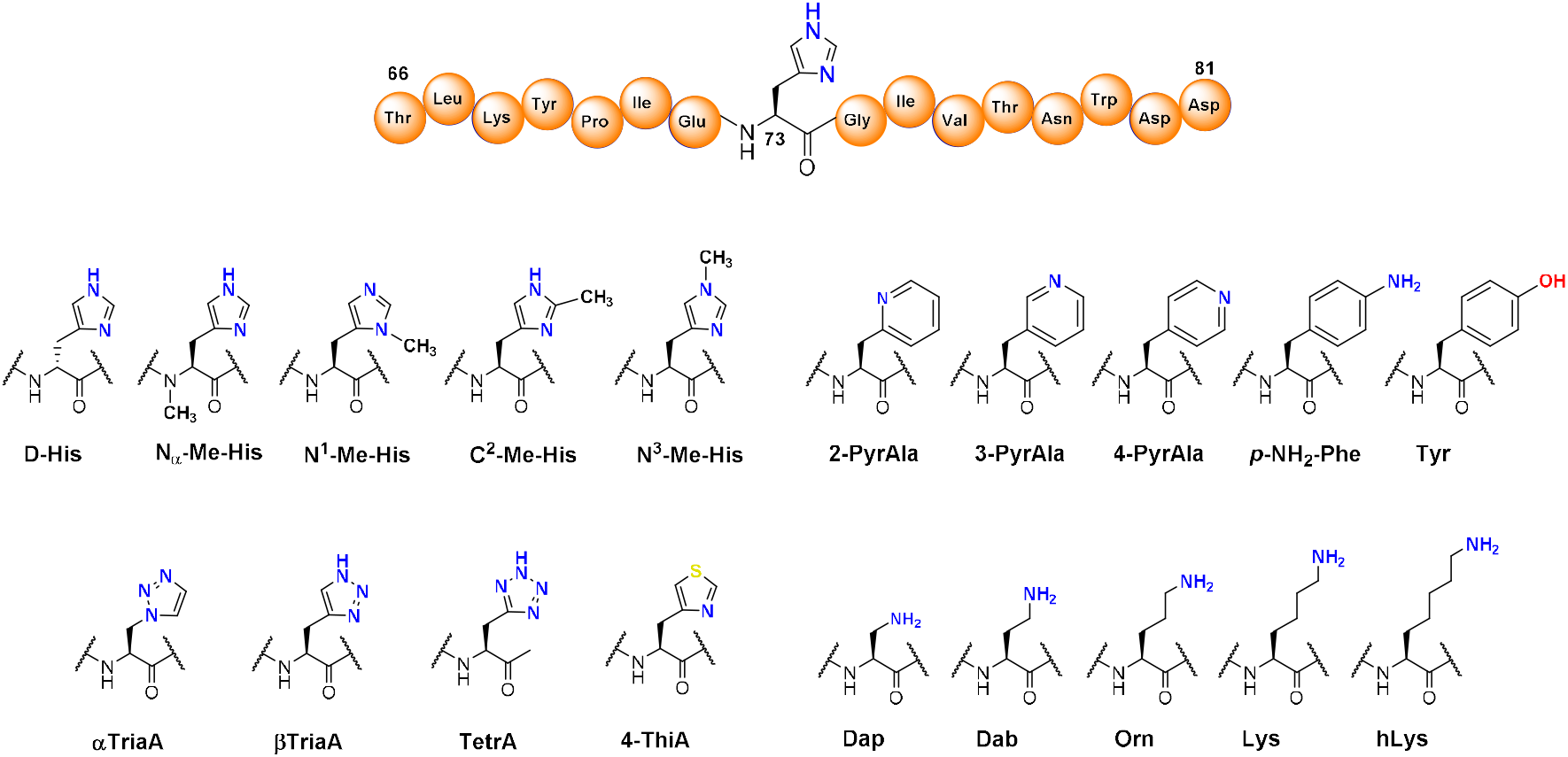
The panel of histidine analogs incorporated in the βA peptide for examination of SETD3 catalysis.

To synthesize the α-triazole, azidoalanine (Aza) was incorporated into the βA peptide directly (Figure S21a). After cleavage from the resin and purification, a biorthogonal copper-catalyzed click reaction enabled the preparation of the target triazole (Siegl et al., 2018). To this end, TMS-acetylene was used, after which the TMS protecting group was removed under acidic conditions to generate the α-triazole. The β-triazole was generated by introduction of propargylglycine (Pra) into the βA peptide and the click reaction was performed using sodium azide, affording the target triazole directly (Figure S21b). C^2^-methyl histidine was generated directly by employing a late-stage functionalization of the histidine-containing βA peptide, using sodium methanesulfinate in the presence of tert-butylhydroperoxide (TBHP) as the oxidant (Figure S21c) (Noisier et al., 2017).

### SETD3-catalyzed methylation by MALDI-TOF MS

We carried out enzymatic assays with the synthetic 16-mer βA peptides using MALDI-TOF MS assays (Hintzen et al., 2021). A βA peptide (10 μM) was incubated in the presence of the recombinantly expressed human SETD3 (1 μM) and SAM cosubstrate (100 μM) at 37 °C (standard conditions). All peptides possessing histidine and its analogs were studied at pH 7.2, pH 9.0 and pH 10.5; enzymatic mixtures were quenched after 1 hour and 3 hours. Under these conditions, the histidine-containing βA-His73 peptide was quantitatively monomethylated in 1 hour at all three pHs, without producing any dimethylated histidine (Figure 3a, Figures S22-23). Control reactions in the absence of either the SAM cosubstrate or the SETD3 enzyme afforded no detectable methylation of the peptide, indicating that methylation reactions are enzymatic and require the SAM cosubstrate (Figure S24). In contrast, D-histidine-containing βA-D-His73 was not shown to be methylated by SETD3 within detection limits, demonstrating that the stereochemistry of the histidine is an important molecular requirement of the substrate methylation (Figure 3b, Figures S22-23). These data are in line with a general lack of enzymatic methylation of D-lysine residues in histones by SET domain histone lysine methyltransferases Belle et al., 2017). Under standard conditions, N_α_-Me-histidine also did not undergo SETD3-catalyzed methylation (Figure 3c, Figures S22-23). We attribute the lack of observed methylation to disruption of an essential H-bond between His73’s main chain amide NH and the Tyr312’s main chain CO in SETD3 that likely contributes to the correct positioning of the His73 side chain for efficient methylation reaction. These results resemble data from enzymatic studies that showed the importance of the lysine’s backbone for efficient KMT-catalyzed methylation reaction (Al Temimi et al., 2019).

**Figure 3.**
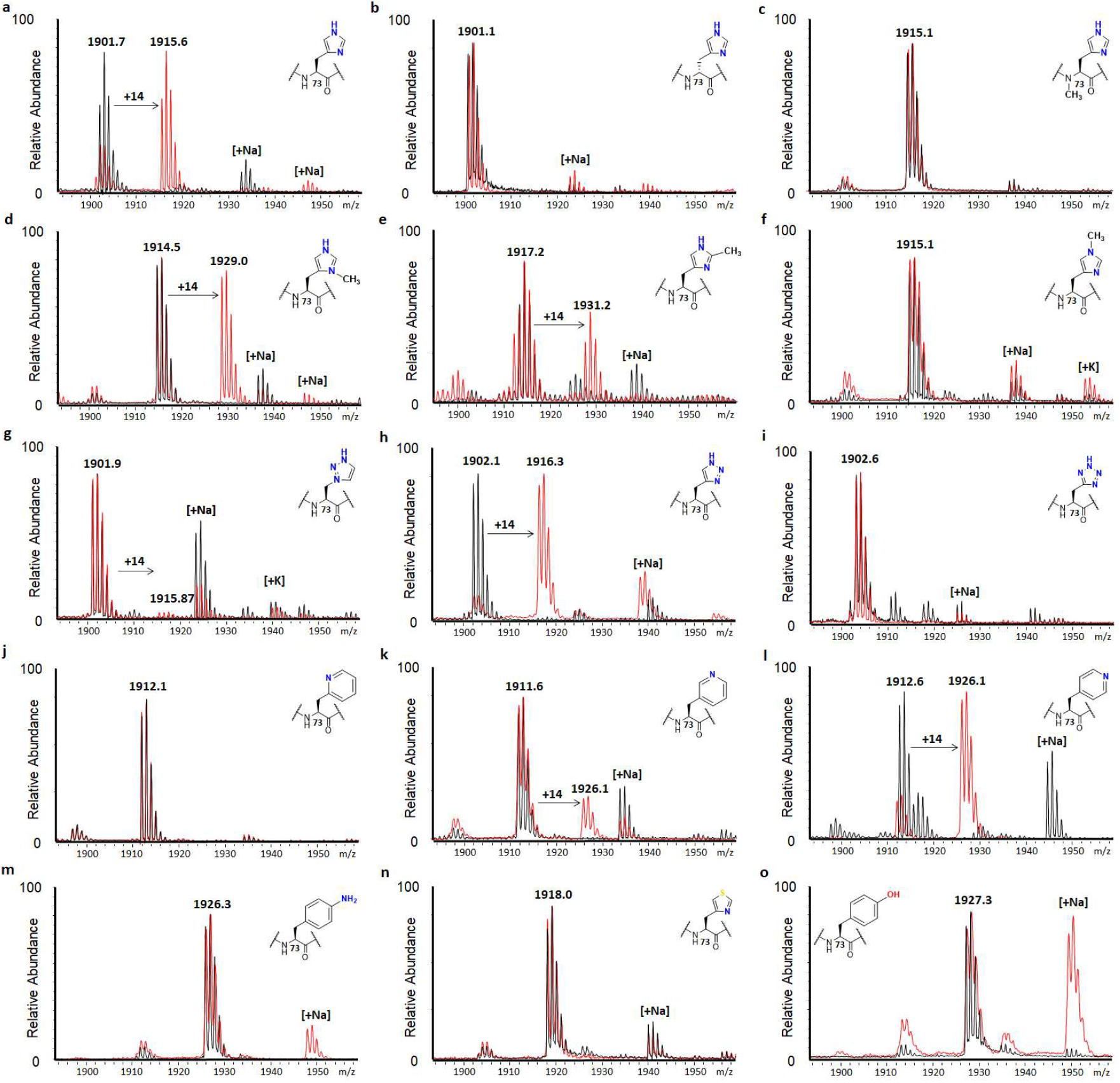
MALDI-TOF MS data showing methylation of βA_66-81_ peptides (10 μM) in presence of SETD3 (1 μM) and SAM (100 μM) after 3 hours at pH 9.0. Control reactions without SETD3 present are shown in black, SETD3-catalyzed reactions are shown in red. **a)** βA-His73, **b)** βA-D-His73, **c)** βA-N_α_-Me-His73, **d)** βA-N^1^-Me-His73, **e)** βA-C^2^-Me-His73, **f)** βA-N^3^-Me-His73, **g)** βA-αTriaA73, **h)** βA-βTriaA73, **i)** βA-TetrA73, **j)** βA-2PyrAla73, **k)** βA-3PyrAla73, **l)** βA-4PyrAla73, **m)** βA-*p*-NH_2_-Phe73, **n)** βA-4ThiA73, **o)** βA-Tyr73.

Interestingly, our MALDI-MS data showed that SETD3 has the ability to catalyze methylation of the premethylated N^1^-Me-His, producing N^1^,N^3^-dimethylhistidine. Previous studies have shown that SETD3 specifically catalyzes methylation at the N^3^-position (Guo et al., 2019), and under our assay conditions trace amounts of dimethylated histidine were found when using the βA-N^1^-Me-His73 substrate at pH 7.2 and 10.5 after 3 hours (Figures S22-23), and 44% of dimethylated product at pH 9.0 after 3 hours (Figure 3d). βA-C^2^-Me-His73 (Figure 3e, Figures S22-23) also underwent SETD3-catalyzed methylation, generating the C^2^,N^3^-dimethylhistidine. In contrast, N^3^-Me-histidine did not undergo SETD3-catalyzed methylation under standard conditions (Figure 3f, Figures S22-23), further cementing SETD3 as a N^3^-specific histidine methyltransferase.

βA-αTriaA73 underwent SETD3-catalyzed methylation to a minor extent after 3 hours at pH 7.2 (Figure S22), while no methylation was observed at pH 9.0 and 10.5 (Figure 3g and Figure S23). Notably, βA-βTriaA73 was found to be fully methylated after 3 hours at pH 7.2 and pH 9.0 (Figure 3h and Figure S22), displaying similarity with the βA-His73 peptide at physiological and slightly basic pH values. At pH 10.5, βA-βTriaA73 was methylated only to a minor extent (Figure S23). This observation could be attributed to the difference in p*K*_a_ values between the imidazole (14.4) and the triazole (9.3), causing the triazole to be predominantly deprotonated at pH 10.5 (Konášová et al., 2015). As the α-triazole does not contain a free hydrogen on a nitrogen atom in the imidazole ring, and the β-triazole does, these results indicate that this hydrogen atom appears to be important for enzymatic activity through the hydrogen bonding stabilization of the ring configuration as observed in the case of βA-His73 (see below). Finally, βA-TetraA73 was not methylated beyond the detection limits at all three pHs (Figure 3i, Figure S22-23). Due to low p*K*_a_ value of tetrazole (4.9), making it a well-known carboxylic acid bioisostere, the tetrazole side chain should be fully deprotonated at pH 7.2, likely leading to nonoptimal binding and the lack of methylation. At lower pH values (3.5 and 5.0), βA-TetraA73 underwent minor methylation in the presence of 1 μM SETD3 (Figure S25) and significant methylation in the presence of 10 μM SETD3 (Figure S26), unlike βA-His73 and βA-βTriaA73 peptides. It has to be noted that all these reactions did not progress between the 1 hour and 3 hour timepoints, indicating that the enzyme itself indeed becomes inactive at low pH (Figure S27). Taken together, these findings seem to support that the electronic properties of the nitrogen-containing aromatic ring are crucial, and that the reactivity of SETD3 can be finetuned across a range of pH values to specifically methylate certain groups.

Of the more sterically demanding six membered rings, none of the pyridylalanine-containing peptides were methylated by SETD3 at pH 7.2 (Figure S22), while strikingly, 3- and 4-pyridylalanine underwent an efficient SETD3-catalyzed methylation at pH 9.0 (Figure 3k,l). Under standard conditions, βA-4PyrAla73 reached 80% conversion after 3 hours, while βA-3PyrAla73 was methylated less efficiently, reaching 20% methylation. At pH 10.5, methylation activity for βA-3PyrAla73 was completely abolished, while βA-4PyrAla73 was still accepted as a substrate relatively well (21% conversion, Figure S24). βA-2PyrAla73, or an exocyclic amino group in βA-*p*-NH_2_-Phe73 were not methylated by SETD3 at all three tested pHs (Figure 3j,m, Figures S23-24). These results suggest that SETD3 is very sensitive to the positioning of the nitrogen atom to be methylated. The orientation in the 4-pyridyl is closest to the original N^3^ of the imidazole ring of His, while the orientation of the 2-pyridyl would line up most with the N^1^ nitrogen of the imidazole, leaving the 3-pyridyl positioned in between the two imidazole nitrogens. SETD3 thus clearly shows a preference for the 4-pyridyl orientation over the 3-pyridyl. Intrigued by the fact that the βA-3PyrAla73 and βA-4PyrAla73 peptides were accepted to a lesser extent at pH 10.5 in comparison to pH 9.0, we investigated whether the methylated product of the reaction is stable under more basic conditions. This revealed that only after 1 hour at pH 10.5, all the methylated 3-pyridyl analog was degraded to 3-pyridylalanine (Figure S28), confirming the finding that no methylation was observed after incubation in reaction buffer with SETD3 at pH 10.5.

Our further examinations revealed that SETD3 catalysis appears to be limited to nitrogen nucleophiles within the scope of our panel. The 4-thiazolyl alanine containing peptide was not methylated, indicating that SETD3 is not capable of methylation of sulfur in an aromatic system, even though it is at the same position as the N^3^ of histidine (Figure 3n, Figures S22-23). Incorporation of tyrosine did not lead to methylation (Figure 3o, Figures S22-23), which could be due to several reasons, including the steric demand of the six membered ring, the existence of an oxygen nucleophile instead of nitrogen, and the target group lying outside of the ring, which was not methylated in the case of an amino group either.

To further investigate the effect of lysine’s aliphatic side chain on the efficiency of the SETD3-catalyzed methylation, βA peptides bearing diaminopropionic acid (Dap), diaminobutyric acid (Dab), ornithine (Orn), lysine (Lys) and homolysine (hLys) were examined under the same assay conditions as described above. In principle, SETD3 might have an ability of catalyzing three subsequent methylation steps on the terminal amino group of the lysine and its analogs, much alike other KMTs. Our results revealed that at pH 7.2 none of the lysine analogs was methylated (Figure S22), while at pH 9.0 methylation was only observed for Dab and Orn to a minor extent (Figure S29). At pH 10.5, however, only minor monomethylation of Dab and Orn was observed (Figure S23). These results indicate that Dab and Orn have chain lengths that are better accommodated by the smaller active site of SETD3, in comparison to SET-domain lysine methyltransferases. However, as previously described (Dai et al., 2019), the lysine containing βA-Lys73 is a much poorer substrate than the histidine containing βA-His73, therefore higher concentrations of enzyme for the panel were used to assess which of lysine analogs possesses the optimal chain length. Lysine analog-containing βA peptides (10 μM) were incubated in the presence of human SETD3 (10 μM) and SAM cosubstrate (200 μM) at 37 °C (Figure 4). Interestingly, only minor monomethylation for Dab (7%) and Orn (14%) was observed already at pH 7.2 after 3 hours, with no lysine methylation observed (Figure S30). At pH 9.0, only Dab, Orn and Lys showed monomethylation, with some dimethylation also observed for Orn (Figure S31). At pH 10.5, similar profiles as at pH 9.0 were observed for Dab and Lys (Figure 4b,d, right panels), however, Orn now showed the major part of the starting material converted, and mono-, di- and trimethylation were readily observed at 35%, 32% and 6% respectively (Figure 4c). Dap and hLys were not found to be methylated under any of the pHs, even in the presence of an increased concentration of SETD3 (Figure 4a,e, Figures S30-31). A consumption curve with βA-Orn73 showed a gradual decrease of the unmethylated βA-Orn73 and a sequential increase of methylated species in a period of 3 hours (Figure 4f).

**Figure 4.**
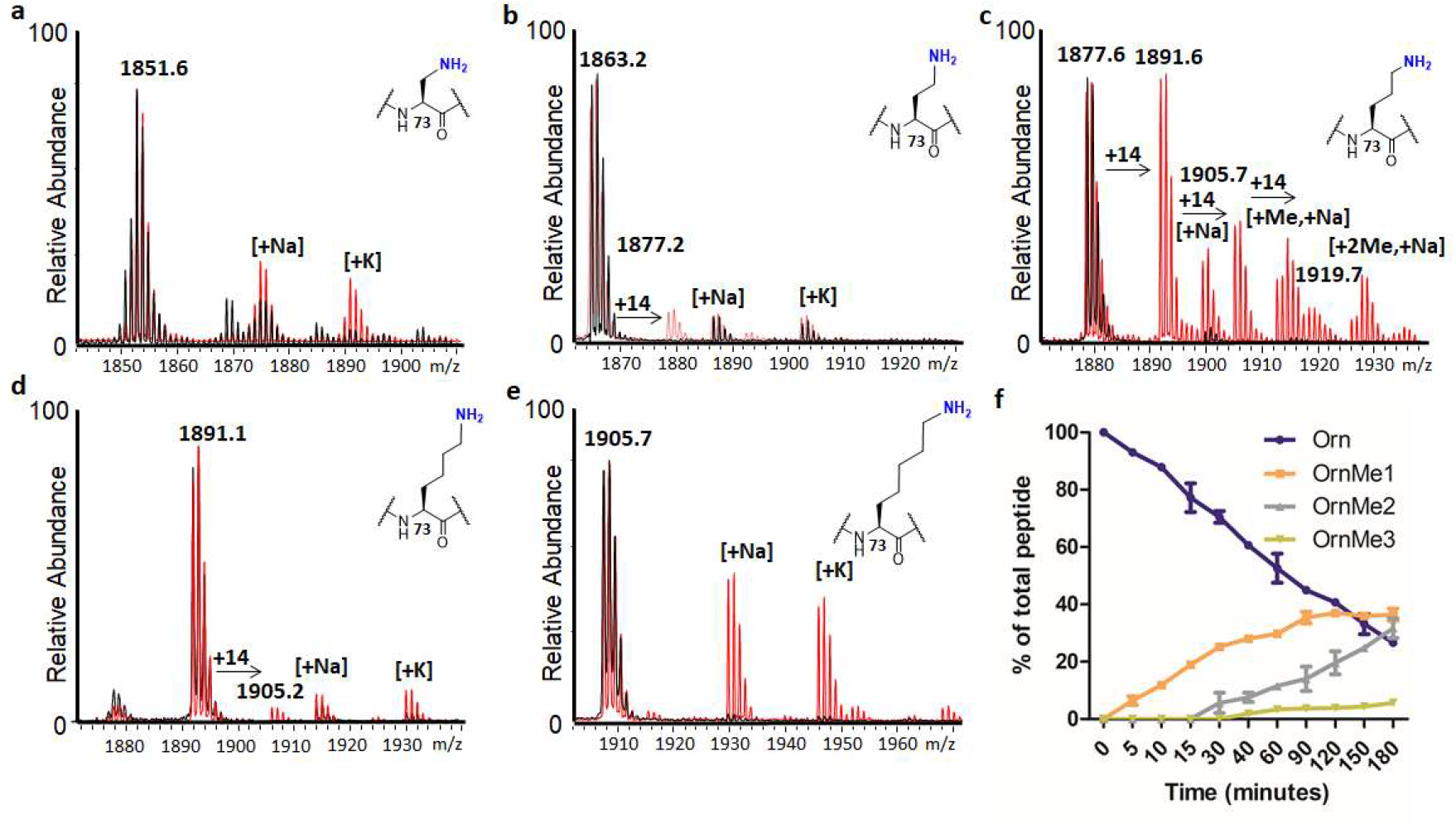
MALDI-TOF MS data showing methylation of βA peptides (10 μM) possessing lysine analogs in presence of increased concentration of SETD3 (10 μM) and SAM (200 μM) after 3 hours at pH 10.5. Control reactions without SETD3 present are shown in black, SETD3 catalyzed reactions shown in red. **a)** βA-Dap73, **b)** βA-Dab73, **c)** βA-Orn73, **d)** βA-Lys73, **e)** βA-hLys73; **f)** Consumption curve of the SETD3-catalyzed methylation of βA-Orn73 at pH 10.5.

### Enzyme kinetics for SETD3-catalyzed methylation of histidine analogs

Having established that some histidine analogs can be accepted for SETD3-catalyzed methylation, we sought to investigate their kinetic profiles. MALDI-TOF MS kinetic studies were used under steady-state conditions varying the concentration of the synthetic βA peptides in the kinetic buffer at pH 9.0. The concentration of SETD3 was adjusted to accommodate linear conversion for the analogs in a time window of 20 minutes at 37 °C. Out of our tested βA peptides, the βA-His73 substrate still had the best catalytic efficiency (k_cat_/K_m_, Table 1, Figure S32a), while βA-4PyrAla73 and βA-βTriaA73 displayed 3-fold and 2-fold decrease in the catalytic efficiency, respectively (Table 1, Figure S32c,e). Interestingly, the catalytic rate k_cat_ for the βTriaA is – within the margin of error – the same as for the natural substrate, the loss in activity here is caused by an apparent loss of binding, as reflected by a higher K_m_ value. On the other hand, it was found that βA-4PyrAla73 and βA-His73 display similar K_m_ values, and the decrease in activity for βA-4PyrAla73 is attributed to a less favorable in k_cat_. These two histidine analogs appeared to be the best substrates within our panel, while the two other tested analogs, βA-N^1^-Me-His73 and βA-3PyrAla73, displayed a significantly lower catalytic efficiency (12- and 6-fold) in comparison to βA-His73 (Table 1, Figure S32b,d). For βA-Orn73 a 25-fold reduced substrate efficiency was observed, however, it was tested at pH 9.0 for direct comparision with other analogs, whereas it was shown to be better accomodated at pH 10.5 (Figure S32f). As suggested from data in Figure 4, and comparing to previously reported results for Lys (Dai et al., 2019) with the SETD3, Orn appears to be a much better substrate than Lys, owing to the fact that ornithine is a better histidine mimic and its side chain can be better positioned into the catalytic pocket of SETD3 for efficient methylation reaction.

**Table 1.** Kinetic parameters for SETD3-catalyzed methylation of βA peptides.

| Peptide | $k_{\text{cat}}(\text{min}^{-1})$ | $K_{\text{m}}(\mu\text{M})$ | $k_{\text{cat}}/K_{\text{m}}$<br>($\mu\text{M}^{-1}\text{min}^{-1}$) |
| --- | --- | --- | --- |
| βA-His73 | $141 \pm 7.4$ | $33.6 \pm 4.7$ | 4.22 |
| βA-N<sup>1</sup>-Me-His73 | $28.2 \pm 2.5$ | $88.6 \pm 16$ | 0.32 |
| βA-βTriaA73 | $138 \pm 9.6$ | $66.7 \pm 11$ | 2.07 |
| βA-3PyrAla73 | $48.8 \pm 3.0$ | $62.9 \pm 9.1$ | 0.78 |
| βA-4PyrAla73 | $47.6 \pm 4.4$ | $33.8 \pm 8.3$ | 1.41 |
| βA-Orn73 | $14.0 \pm 1.2$ | $95.8 \pm 16$ | 0.15 |

Next, we tested the analogs that were not accepted for SETD3-catalyzed methylation for their potency as inhibitors of SETD3. Due to difficulties with overlapping signals in our mass spectrometric assay, βA-N_α_-Me-His73, βA-N^1^-Me-His73, βA-αTriaA73 and βA-TetrA73 were omitted from the panel, whereas analogs that were not methylated were tested in a single-point screening assay. SETD3 (180 nM) was incubated for 1 hour at pH 9.0 with the potential inhibitory peptides (100 μM) and SAM (100 μM), subsequently the βA-His73 peptide (10 μM) was added and reacted for 20 minutes to ensure linear conversion to the methylated species. All analogs containing an aromatic ring-structure in the side chain, along with the lysine analogs Dap and hLys were found to be inactive, with apparent IC_50_ values above 100 μM (Figure S33).

### Calorimetric titration studies of SETD3 with analog βA peptides

To explore the substrate specificity of SETD3, we measured the binding affinities by isothermal titration calorimetry (ITC) of SETD3 with His73 or selected histidine analog-replaced βA peptides in the presence of SAH and sinefungin (SFG), respectively. ITC titration experiments showed that the binding *K*_D_ values dropped from 0.7 μM for wild type βA-His73 to 0.8, 3.1, 7.2, and 44.4 μM for the βA-βTriaA73, βA-4PyrAla73, βA-Orn73, and βA-3PyrAla73 peptides in the presence of SAH (Figure 5c, Figures S34-41). Corresponding *K*_D_ values are 1.0, 4.0, 4.9, 39.2, 48.5 μM in the presence of SFG (Figure 5d). The measured binding affinities in the presence of SAH are generally stronger than those in the presence of SFG, consistent with the previous report, and suggest the existence of steric tension between the bulkier SFG with the βA peptides at the active center (Zheng et al., 2020).

**Figure 5.**
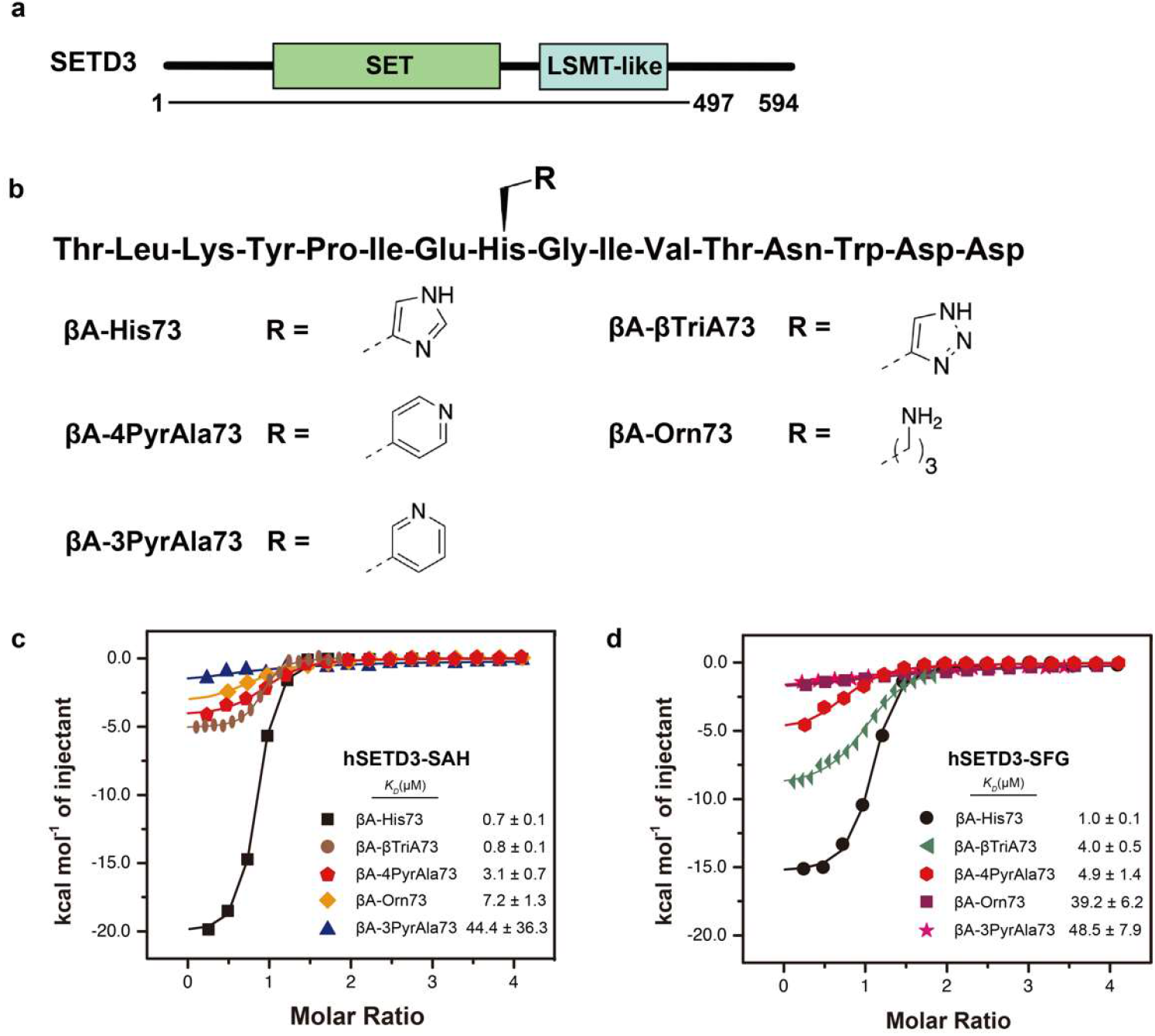
Binding studies of SETD3 with the βA peptide and its His73-replaced analogs. **a)** Domain architecture of human SETD3. Underlined region is the expression frame of SETD3 for binding study. **b)** The sequence and functional group information of peptides used for titration and crystallization. **c, d)** ITC fitting curves of SETD3_1-497_ titrated with wild type β-actin peptide and analogs mentioned above in the presence of SAH or SFG.

### Co-crystal structural studies of SETD3 bound to analog βA peptides

To elucidate the molecular basis for substrate engagement by SETD3, we determined the crystal structures of SETD3-βA-Orn73-SAH (PDB: 7W29) and SETD3-βA-4PyrAla73-SAH (PDB: 7W28) ternary complexes at 2.9 Å and 1.8 Å resolution, respectively (Table S2). The overall engagement modes of histidine analogs with SETD3-SAH were similar to that of the native βA peptide. In the complex structure, the His73 analog and SAH are confined in the catalytic channel of SETD3 through a “head-to-head” mode (Figure 6a and b, close-up view). The peptides of βA-4PyrAla73 and βA-Orn73 can be clearly traced and modelled according to the 2Fo-Fc omit map (Figure 6c). Superimposition of the wild type (PDB: 6MBJ) and the βA-Orn73 ternary complex structures revealed similar substrate engagement modes. We observed similar positioning (∼0.6 Å shift) of the nitrogen atom of Orn73 terminal amine and the N^3^ atom of histidine next to the sulfur atom of SAH at the active center, consistent with the decent methyltransferase activity measured for the βA-Orn73 substrate (Figure 6d). In the meantime, we observed clear twist of the C_α_ backbone of Orn73 as compared to His73 of wild type βA. This suggested steric tension likely caused by the bulkier size of the Orn73 side chain, thus accounting for the weaker K_m_ or *K*_D_ observed for the βA-Orn73 analog. Collectively, these results highlight exquisite substrate recognition mechanisms that may determine the substrate preference of SETD3. Structural alignment of the wild type βA-His73-SAH (PDB: 6MBJ) and the βA-4PyrAla73-SAH ternary complexes displayed the similar positioning of the N^3^ atom of histidine and the nitrogen atom of 4PyrAla73 pyridine ring at the active center, consistent with the observed enzymatic activity (Figure 6e). Meanwhile, the bulkier size of the six-atom ring of 4PyrAla73 side chain may lead to four-fold decease of the binding affinity in the presence of SAH (Figure 5). Further superimposition of SETD3-βA-4PyrAla73-SAH, SETD3-βA-His73-SAH and SETD3-βA-His73-SFG (PDB: 6JAT) revealed rotational adjustments of the rings at residue 73 to better fit the dimension of the catalytic pocket (Figure 6f). Collectively, our structural comparison analyses support conformational flexibility at the active center, which is responsible for the observed substrate diversity in this study.

**Figure 6.**
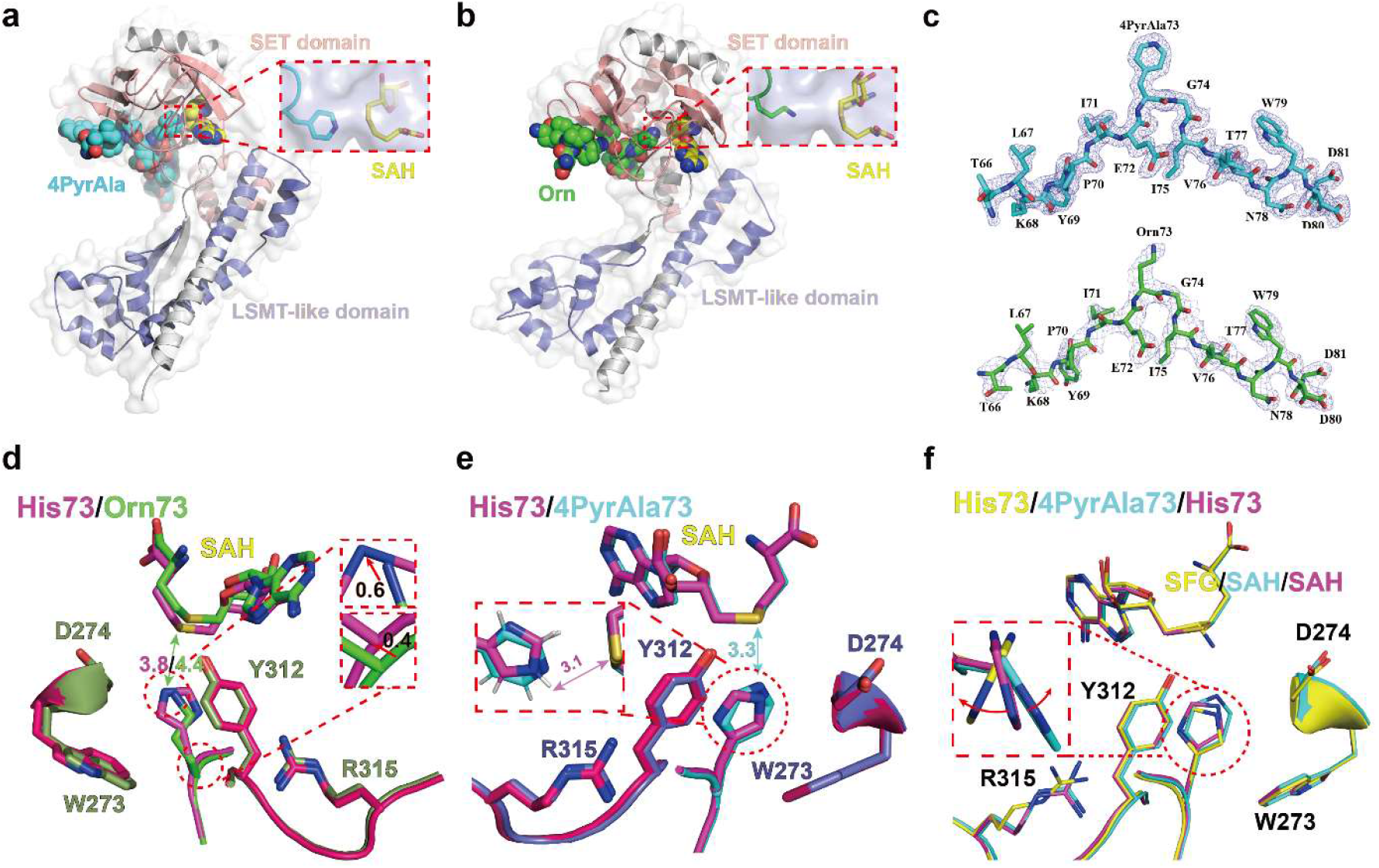
Structural characterization of SETD3 bound to histidine analog βA peptides. **a, b)** Overall structures of SETD3 bound to βA-4PyrAla73 or βA-Orn73 peptide and SAH. The SET and LSMT-like domains of SETD3 are color-coded as indicated. SETD3 is shown in semitransparent surface view. 4PyrAla (cyan) or Orn (green) replaced in the βA peptides and SAH (yellow) are depicted as spheres, respectively. Close-up views, positioning of SAH and the methyl acceptor residues inside the catalytic channel. **c)** 2Fo-Fc omit map of βA-4PyrAla73 and βA-Orn73 peptides contoured at 1.0 σ level. **d**,**e)** Interaction details of peptidyl substrates inserted into the active center of SETD3. His73 (red), Orn73 (green), and 4PyrAla73 (cyan) are shown as sticks, and distances between indicated atoms are labelled in the unit of angstrom with coded color. Functional groups of Orn73 and His73 are also depicted as color-coded meshes in close-up view of panel d. Key residues of SETD3 are depicted as green (Orn73) and blue (4PyrAla73) sticks. **f)** Superimposition of SETD3 in complex with βA-4PyrAla73 (cyan) and βA-actin in the presence of SAH (red) or SFG (yellow).

### QM/MM MD and free energy simulations

Computer simulations were performed on βA-His73, βA-N^1^-Me-His73, βA-βTriaA73, βA-3PyrAla73 and βA-4PyrAla73 to provide additional information concerning the ability of SETD3 to catalyze methylation of these analogs as well as the energetic and structural origins of the activities. The crystal structure complex containing the βA-4PyrAla73 peptide obtained in this work was used to build the models (see methods). The free energy profiles for methylation of βA-His73 and its mimics are plotted in Figure 7. As can be seen from Figure 7, the βA-His73 and βA-βTriaA73 substrates have the lowest free energy barrier (15.4 and 17.4 kcal mol^-1^, respectively). The results are consistent with the kinetics data in Table 1 showing that SETD3 is the most active on βA-His73 and βA-βTriaA73 with the two highest k_cat_ values. For βA-N^1^-Me-His73, βA-3PyrAla73, βA-4PyrAla73, the free energy barriers are relatively higher, and these results are also consistent with the lower k_cat_ values observed for these analogs.

**Figure 7.**
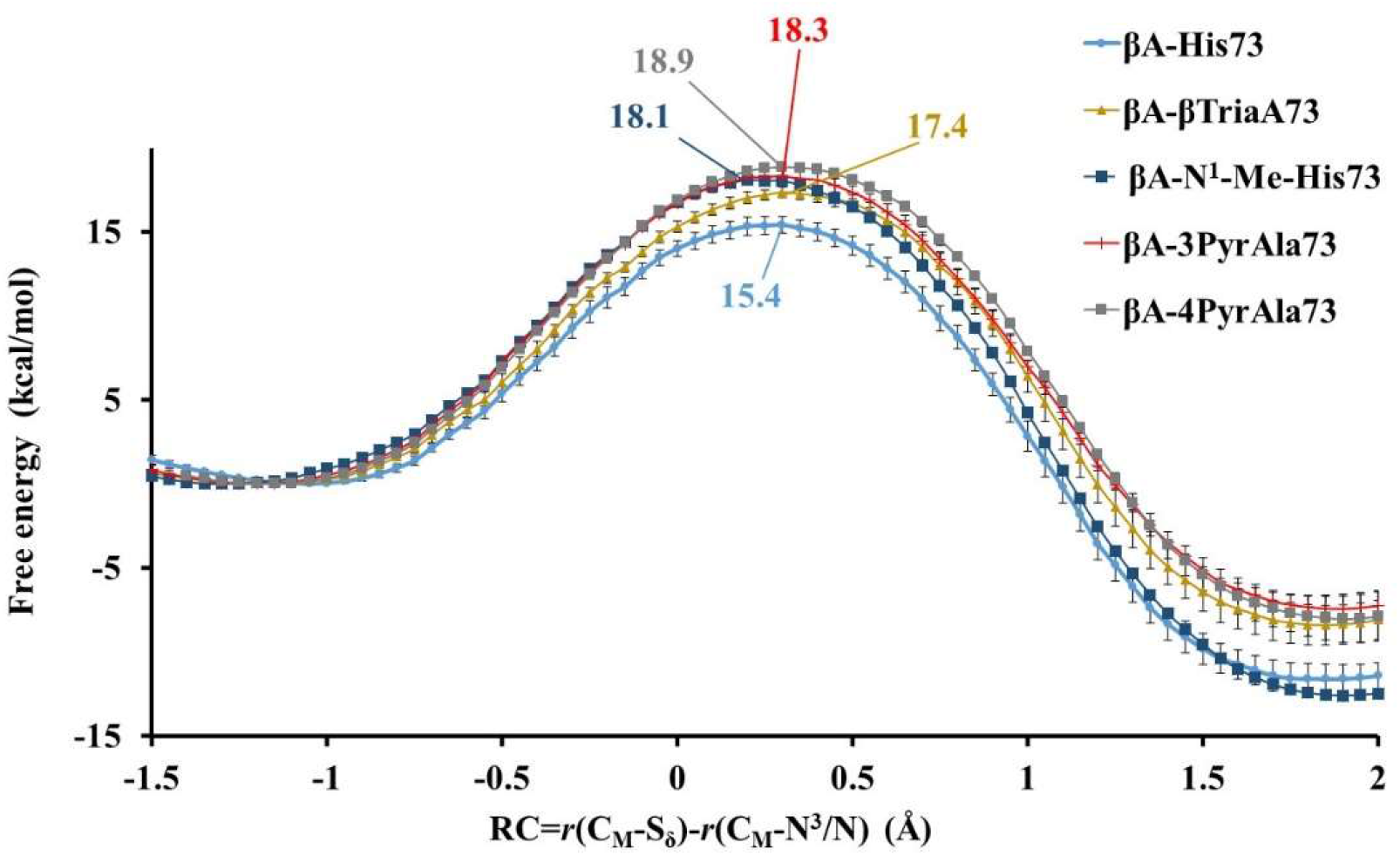
Free energy (PMF) profiles for the methyl transfer from SAM to the target N^3^/N for βA-His73, βA-N^1^-Me-His73, βA-βTriaA73, βA-3PyrAla73 and βA-4PyrAla73, respectively, as a function of the reaction coordinate [R = r(C_M_-S_δ_) -r(C_M_-N^3^/N)] obtained from the QM/MM free energy simulations.

The average structures of the reactive state prior to the methyl transfer are given in Figure 8; other structural information obtained from the simulations are given in Figures S42-47. Figure 8a shows that the structure for βA-His73 is quite close to the one obtained earlier based on a different crystal structure. For instance, there is a hydrogen bond between N^1^-H and Asn255, which stabilizes an orientation of the imidazole ring similar to that in the crystal structures of the product complex (Guo et al., 2019; Dai et al., 2019; Zheng et al., 2020; Deng et al., 2020). It is of interest to notice that this hydrogen bond involving Asn255 also exists in the βA-βTriaA73 complex (Figure 8b). The existence of this hydrogen bond is expected to play an important role in stabilizing the configuration of the βTriaA73 ring observed in the reactive state. The target N^3^ seems to be better positioned to accept the methyl group from SAM at such configuration (Figure 8b), and this may lead to a higher activity of SETD3 on βA-βTriaA73 as observed experimentally. Consistent with this suggestion, previous experimental (Guo et al., 2019, Dai et al., 2019) and computational studies (Deng et al., 2020) on βA-His73 have shown that the k_cat_ (k_cat_/K_M_) value decreased and the free energy barrier increased as a result of the Asn255Ala mutation, which removed the hydrogen bond involving Asn255. Figure 8c and 8d show that the rings of 3PyrAla73 and 4PyrAla73 underwent some rotations relative to those observed for βA-His73 and βA-βTriaA73 with χ(N^1^/C_δ1_−C_β_−C_γ_−C_α_) increasing about 40-70°. The alignments of the transferable methyl group on SAM with the lone-pair electron on N seem not as good as those observed for βA-His73 and βA-βTriaA73, and this may be related to the observations that SETD3 is less active on either of these two analogs (Table 1).

**Figure 8.**
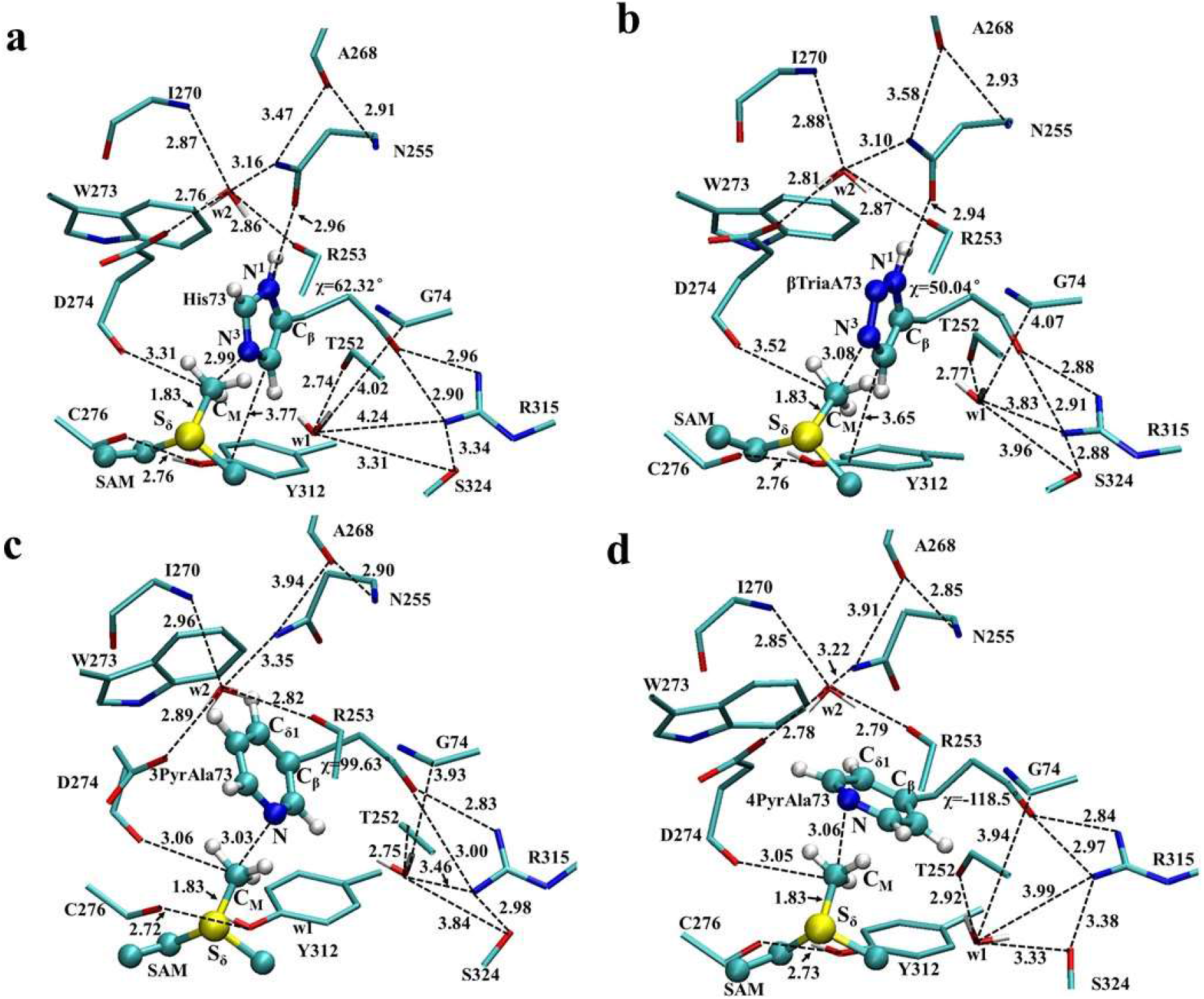
The average active-site structures of the reactive state obtained from QM/MM MD simulations. **a)** βA-His73, **b)** βA-βTriaA73, **c)** βA-3PyrAla73, **d)** βA-4PyrAla73. The average structure for βA-N^1^-Me-His73 and other structural information (e.g., those near transition state) are given in the Supporting Information.

### Conclusion

Understanding protein posttranslational modifications is of central importance to structure and function of proteins, most notably established for histone proteins and their regulation of eukaryotic transcription (Walsh, 2006; Müller and Muir, 2015; Strahl and Allis, 200). Histidine is a subject of several types of PTMs, including N^1^- and N^3^-methylation (Guo et al., 2019, Dai et al., 2019, Davydova et al., 2021), N^1^- and N^3^-phosphorylation (Potel et al., 2018; Fuhs and Hunter, 2017), and C_β_-hydroxylation (Ge et al., 2012; Bundred et al. 2018), most of them being only recently identified and characterized. To advance basic molecular understanding of histidine PTMs, it is essential to explore whether simplest histidine mimics are tolerated and modified by enzymes that install PTMs on histidine residues in proteins. The amber suppression method enabled incorporation of histidine mimics into proteins, however, it is limited to only few selected histidine analogs (Sharma et al. 2016, Xiao et al., 2014). While new chemical tools that rely on click-based phosphohistidine have enabled deeper biomolecular understanding of histone phosphorylation in the context of protein science (Kalagiri et al. 2021; Kee et al., 2010; Makwana et al., 2018), such and related tools for histidine methylation are currently lacking, leading to limited understanding of biologically important histidine methylation. To develop new tool compounds and deepen fundamental understanding of enzymatic histidine methylation, we have here for the first time explored a large panel of histidine mimics as potential substrates for methyltransferase SETD3, establishing its substrate specificity and biocatalytic potential. Our coordinated synthetic, enzymatic, biostructural and computational studies provide strong evidence that human SETD3 has broader substrate specificity beyond histidine. Among the unnatural amino acids in our panel, β-triazolylalanine and 4-pyridinylalanine were observed as excellent SETD3 substrates, whereas N^1^-methylhistidine, 3-pyridinylalanine, ornithine were good, but comparatively poorer substrates. All five histidine analogs, however, display superior substrate efficiency relative to lysine and methionine, two natural amino acids that were recently identified as SETD3 substrates when incorporated into actin peptide and protein (Dai et al., 2020a; Dai et al.; 2020b). Structural analyses of the ternary complexes suggest that SETD3 is a methyl acceptor geometry-sensitive histidine methyltransferase because of its “head-to-head” engagement mode at the catalytic center. The substrate specificity of SETD3 is largely affected by the size compatibility between the acceptor group and the methyl donor SAM. Quantum mechanics/molecular mechanics molecular dynamics and free energy simulations further supported our experimental findings by revealing that some of these mimics could undergo methylation, and the efficiency of the methylation seems to correlate with the ability of the analogs to form the near attack conformations at the active site. Our work, together with recent studies on lysine and methionine methylation, demonstrates that SETD3’s unique ability to catalyze methylation of histidine residues can be extended to efficient methylation of its simple mimics. More generally, this work shows that a combination of state-of-the-art experimental and computational approaches enables investigations on enzymatic catalysis with an unprecedented level of molecular detail.

## Materials and Methods

### SETD3 expression and purification

The recombinant human SETD3 fused to an N-terminal His_6_-tag was produced and purified by a slightly modified procedure described by Kwiatkowski et al., 2018. Briefly, the enzyme was expressed in *E. coli* BL21(DE3) at 13°C overnight in the presence of 0.3 mM IPTG. The recombinant protein was purified with HisTrap FF column (5 ml) and eluted in 20 ml of 50 mM HEPES pH 7.5, 400 mM NaCl, 10 mM KCl, 300 mM imidazole, 1 mM DTT. The elution buffer was exchanged by sequential dialysis of the enzyme preparation against 500 ml of dialysis buffer (20 mM Tric-HCl pH 7.5, 200 mM NaCl, 1 mM DTT and 6% sucrose) overnight at 4 °C and then twice against 500 ml of the buffer for 3h at RT. The enzyme purity was verified by SDS-PAGE (>97%). The purified enzyme was aliquoted and stored at -70°C.

### Synthesis and purification of β-actin peptides

All β-actin peptides were chain assembled on Rink amide resin using microwave assisted SPPS on a Liberty Blue peptide synthesizer (CEM corporation, Matthews, NC, USA). All amino acid couplings were carried out with the equivalent ratio of [5]:[5]:[7.5] of [Fmoc-protected amino acid]:[DIC]:[Oxyma Pure] at 75 °C for 2 minutes or at 50 °C for 4 minutes for histidine, histidine analogs were coupled at 50 °C for 10 minutes. Peptides proceeded to standard cleavage from resin using 0.5% TIPS, 0.5% H_2_O in conc. TFA. The azide containing peptide was cleaved with 0.5% TIPS, 0.5% H_2_O, 0.5% TES and 0.5% thioanisole in conc. TFA. TFA was blown off using N_2_ and the resultant residue suspended in cold Et_2_O. After suspension it was subjected to centrifugation for 5 minutes at 5000 rpm in an Eppendorf 5804R centrifuge (Eppendorf, Hamburg, Germany) after which the supernatant was decanted into the waste. The remaining white to yellow solid was washed twice by cold Et_2_O and subjected to centrifugation after which the crude peptide was dissolved in a mixture of ACN in H_2_O and purified using preparative reverse-phase HPLC (RP-HPLC) using a gradient of buffer A and buffer B from 20% B to 70% over 40 minutes at 4 ml/min using a Gemini 10μm NX-C18 110Å LC column (Phenomenex, Torrance, CA, USA). Analytical RP-HPLC was carried out on a Gemini 5μm C18 110Å LC column (Phenomenex) at a flow rate of 1 mL/min. Analytical injections were monitored at 215 nm.

### General strategy for the on-resin generation of triazolyl alanines

The click reagent (NaN_3_ or TMS-acetylene, 45 eq.) was dissolved in 300 μL MQ and then added to the βA peptide resin under argon atmosphere, followed by short mixing on vortex. CuSO_4_ (6 eq.) was then dissolved in 200 μL MQ and mixed with BTTP (2.6 eq.) followed by addition of sodium ascorbate (4 eq,) dissolved in 200 μL MQ under argon atmosphere. This mixture was added to the peptide solution, followed by the addition of 40 μL DIPEA, and was then reacted overnight at RT while shaking. The resin was then washed with DMF and DCM and subsequently TFA cleaved under standard conditions, diluted with 300 μL ACN and directly purified on RP-HPLC using a gradient of buffer A and buffer B from 20% B to 70% over 40 minutes at 4 ml/min and subsequently lyophilized.

### Synthesis of C^2^-Me-His

The βA peptide was dissolved in H_2_O:DMSO 9:1 (v/v) and heated to 70 °C under argon atmosphere. A solution of sodium methanesulfinate (12 eq., 0.6 M) and TBHP (20 eq., 0.1 M) was added dropwise using a syringe pump (flowrate 100 μL/min) and the reaction was left to stir overnight. The mixture was then diluted with ACN to a ratio of H_2_O:ACN:DMSO 4.5:5:0.5 (v/v/v) and directly purified on RP-HPLC using a gradient of buffer A and buffer B from 20% B to 70% over 40 minutes at 4 ml/min and subsequently lyophilized (Noisier et al., 2017).

### MALDI-TOF MS enzymatic assays

SETD3’s enzymatic activity towards βA peptides was measured at different time points under standard conditions (1 μM SETD3 enzyme, 10 μM β-actin peptide, 100 μM SAM cosubstrate) in the reaction buffer at five different pH’s: 3.5 (21.3 mM sodium citrate dihydrate, 78.7 mM citric acid, 20 mM NaCl), 5.0 (57.7 mM sodium citrate dihyrdate, 42.3 mM citric acid, 20 mM NaCl), 7.2 (25 mM Tris-HCl, 20 mM NaCl), 9.0 (25 mM Tris-HCl, 20 mM NaCl) and 10.5 (20 mM glycine, 20 mM NaOH, 50 mM NaCl). The reactions were carried out in a final volume of 50 μL by incubation by shaking in a Thermomixer C (Eppendorf) at 750 rpm, at 37°C. Lysine analogs and experiments carried out at pH 3.5 and 5.0 were incubated with elevated SETD3 concentrations as well (10 μM SETD3, 10 μM peptide, 200 μM SAM); these reactions were carried out in a final volume of 25 μL. All reactions were quenched by the addition of 10% TFA in MilliQ water (v/v), aliquoted and mixed 1:1 with α-Cyano-4-hydroxycinnamic acid (CHCCA) matrix dissolved in a mixture of H_2_O and ACN (1:1, v/v), and loaded onto an MTP 384 polished steel target to be analyzed by a UltrafleXtreme-II tandem mass spectrometer (Bruker).

### MALDI-TOF MS kinetic assay

βA peptide kinetic evaluation was carried out with a MALDI-TOF MS assay under steady-state conditions at pH 9.0. βA peptides (0-125 μM) were incubated and reactions were started by addition of SETD3 (100-400 nM, depending on the tested analog) in a final volume of 25 μL. The reactions were carried out by incubation by shaking in a Thermomixer C (Eppendorf) at 750 rpm, at 37°C. Steady-state conditions were guaranteed by saturating concentrations of SAM (1 mM, >5x K_m_ value). Reactions were quenched by the addition of 10% TFA in MQ after 20 minutes. All reactions were aliquoted, mixed 1:1 with α-Cyano-4-hydroxycinnamic acid (CHCCA) in a mixture of H_2_O and ACN (1:1, v/v) and loaded onto an MTP 384 polished steel target to be analyzed by a MALDI-TOF MS. The amount of methylated peptide was calculated by intergation of the product peak area and divided by the amount of total (methylated and unmethylated) peptide, taking in account all the ionic species. Kinetic values were extrapolated by fitting V_0_ values and βA peptide concentrations to the Michealis-Menten eqution using GraphPad Prism 5. Experiments were carried out in duplicate and final values are reported as value ± SD.

### MALDI-TOF MS inhibition studies

βA peptides (10 μM), SAM (100 μM) and SETD3 (180 nM) were preincubated for 1 hour at 37°C at pH 9. Then, the natural βA peptide (sequence 66-81) (10 μM) was added to the mixture to a final volume of 50 μL and incubated for an additional 20 minutes at 37°C. The reactions were quenched by the addition of TFA 10 % in MilliQ water, aliquoted and mixed 1:1 with α-Cyano-4-hydroxycinnamic acid (CHCCA) matrix dissolved in a mixture of H_2_O and ACN (1:1, v/v), and loaded onto an MTP 384 polished steel target to be analyzed. SETD3 residual activity was determined by calculating the relative integral of the methylated peptide to a control reaction in absence of potential inhibitory peptides. Experiments were carried out in duplicate.

### Isothermal titration calorimetry (ITC)

The titration was performed using the MicroCal PEAQ-ITC instrument (Malvern Instrument) at 25°C. Before ITC titration, SETD3 was incubated with SAH or SFG in a molar ratio of 1:10 for 1 hour in buffer 20 mM Tris-HCl, pH 8.0 and 150 mM NaCl. Each ITC titration consisted of 17 successive injections. Usually, peptides at 0.5-1 mM were titrated into the complex of SETD3 and SAH or SFG at 0.05 mM. The resultant ITC curves were analyzed with Origin 7.0 (OriginLab) using the “One Set of Binding Sites” fitting model. Protein concentrations were measured based on the UV absorption at 280 nm. The dissociation constants (K_d_s) were determined from a minimum of two experiments (mean ± SD).

### Crystallization, data collection, and structural determination

Crystallization was performed via the sitting drop vapor diffusion method at 16°C by mixing equal volume (0.2-1μl) of protein with reservoir solution. SETD3_1-497_ was premixed with SAH in a 1:10 molar ratio in buffer containing 20 mM Tris-HCl, pH 8.0 and 150 mM NaCl. The protein sample was prepared by mixing SETD3_1-497_-SAH complex with peptides in a molar ratio of 1:3 overnight at 4°C. The crystal of SETD3_1-497_-βA-4PyrAla73-SAH was grown in a reservoir solution containing 30%(w/v) PEG4000, 0.1 M Tris-HCl, pH 8.5, 0.2 M Li_2_SO_4_. And the reservoir solution of SETD3_1-497_-βA-Orn73-SAH was 20%(w/v) PEG6000, 0.1 M Bicine-NaOH, pH 9.0. The co-crystals were briefly soaked in cryoprotectant, composed of reservoir solution supplemented with 10% glycerol, and then flash-frozen in liquid nitrogen for data collection. Diffraction data of SETD3_1-497_-βA-Orn73-SAH was collected at beamline BL17U1 at Shanghai Synchrotron Radiation Facility at 0.9792 Å. And the diffraction data set of SETD3_1-497_-βA-4PyrAla73-SAH was collected at beamline BL19U2 at 0.9788 Å. Diffraction images were indexed, integrated, and merged using the HKL2000 software (Otwinowski and Minor, 1997). The structures were solved by molecular replacement using Molrep (Vagin and Teplyakov, 2010) from the CCP4 suite with SETD3-βA-His73-SAH (PDB ID: 6MBJ) as the search model. Refinement and model building was performed with PHENIX (Adams et al., 2010) and COOT (Emsley and Cowtan, 2004), respectively. The data collection and structural refinement statistics are summarized in Table S2.

### Computational Analysis

QM/MM MD and free energy (PMF) simulations were performed to study the structural properties and dynamics of the reactive state of the enzyme-substrate complex for methylation and to calculate the free energy profile for the methyl transfer from SAM to N^1^/N of the target His73 or analog (βA-His73, βA-N^1^-Me-His73, βA-βTriaA73, βA-3PyrAla73 or βA-4PyrAla73) in wild-type SETD3 using the CHARMM program (Brooks et al., 1983). The – CH_2_–CH_2_–S^+^(Me) –CH_2_– part of SAM and the analog sidechain were treated by QM and the rest of the system by MM. The link-atom approach (Field et al., 1990) was applied to separate the QM and MM regions. A modified TIP3P water model (Jorgensen et al., 1983) was employed for the solvent, and the stochastic boundary molecular dynamics method (Brooks et al., 1985) was used for the QM/MM MD and free energy simulations. The system was separated into a reaction zone and a reservoir region, and the reaction zone was further divided into a reaction region and a buffer region. The reaction region was a sphere with radius *r* of 20 Å, and the buffer region extended over 20 Å ≤ *r* ≤ 22 Å. The reference center for partitioning the system was chosen to be the N_ε2_ atom of the His residue. The resulting systems contained around 5700atoms, including about 500 water molecules. The DFTB3 method (Elstner et al., 1998; Cui et al., 2001; Christensen et al., 2016; Lu et al.; 2015) implemented in CHARMM was used for the QM atoms and the all-hydrogen CHARMM potential function (PARAM27) (MacKerell et al., 1998) was used for the MM atoms. The initial structures for the entire stochastic boundary systems were optimized using the steepest descent (SD) and adopted-basis Newton-Raphson (ABNR) methods. The systems were gradually heated from 50.0 to 298.15 K in 50 ps. A 1-fs time step was used for integration of the equation of motion, and the coordinates were saved every 50 fs for analyses. 5ns QM/MM MD simulations were carried out for each of the structures of the reactive state and the distribution maps of *r*(C_M_-N_ε2_) and *θ* were generated in each case; here *θ* is defined as the angle between the direction of the C_M_-S_δ_ bond and the direction of the electron lone pair on N_ε2_. Classical MD simulations (30 ns) with CHARMM force field were also performed to confirm the structure and stability of the reactive state for methylation. To make sure that our computational approaches can re-produce the experimental structures, the QM/MM MD simulations were also performed for the product complex based on the crystal structures. It was found that the experimental structures could be well re-produced using our computational approaches.

The structures of the reactive state for methylation were generated based on the crystal structure of the SETD3 complex containing the βA-4PyrAla73 peptide obtained in this work. SAH in the crystal structure was changed to SAM manually and the βA-4PyrAla73 peptide was changed to βA-His73, βA-N^1^-Me-His73, βA-βTriaA73 or βA-3PyrAla73 to study the methylation process involving each of these peptides; for βA-4PyrAla73, the coordinates from the crystal structure were used directly. The imidazole ring of His73 was generated as the N^1^-H π tautomer with N^3^ unprotonated and N^1^ protonated before the simulations were performed; the imidazole ring rotated around the C_β_-C_γ_ bond during the QM/MM MD simulations, leading to the orientation similar to that in the product complex (Guo et al., 2019; Dai et al., 2019). The umbrella sampling method (Torrie and Valleau, 1974) implemented in the CHARMM program along with the Weighted Histogram Analysis Method (WHAM) (Kumar et al., 1992) was applied to determine the change of the free energy (potential of mean force) as a function of the reaction coordinate for the methyl transfer from SAM to His73/analog in SETD3. The reaction coordinate was defined as a linear combination of *r*(C_M_-N^3^/N) and *r*(C_M_-S_δ_) [*R* = *r*(C_M_-S_δ_)– *r*(C_M_-N^3^/N)]. For the methyl transfer process, 25 windows were used, and for each window 100 ps production runs were performed after 50 ps equilibration. The force constants of the harmonic biasing potentials used in the PMF simulations were 50 to 400 kcal mol^−1^ Å^−2^. In each case, five independent PMF simulations were performed. The free energies (PMFs) and statistical errors were taken as the average values and standard deviations from the five runs, respectively. High-level ab initio calculations (i.e., B3LYP/6-31G**) are too time-consuming for generating free energy profiles of enzyme-catalyzed reactions. The semi-empirical approach based on DFTB3 has been used previously on a number of systems, and the results seem to be quite reasonable for the estimate of the relative free energy barriers along a given reaction coordinate involving wild-type and mutants (Christensen et al., 2016; Lu et al., 2015). It should be pointed out that the bond breaking and making events studied in this work involve simple and similar S_N_2 methyl transfer processes so that much of the errors for the methyl transfers in His73 and different analogs are presumably similar and cancelled out when the relative free energy profiles are compared.

### Accession codes

Coordinates of SETD3-βA-4PyrAla73 and SETD3-βA-Orn73 have been deposited into the Protein Data Bank under accession codes 7W28 and 7W29, respectively.

## Supporting information

Experimental and computational details, characterization of peptides, enzymatic supporting figures, crystallographic supporting information, computational supporting figures.

## Author Information

### Notes

The authors declare no competing interests.

## Acknowledgements

This research was supported by the European Research Council (ERC Starting Grant, ChemEpigen-715691 to J.M.), an Opus-14 grant from the National Science Centre, Poland (2017/27/B/NZ1/00161 to J.D.), Natural Science Foundation of China (22177064 to P.Q.), the Natural Science Foundation of Shandong Province (ZR2021MB050 to P.Q.), National Natural Science Foundation of China (31725014 to H.L.) and National Key Research Development Program of China (2020YFA0803300 to H.L.).

